# Microalgal pigments and their relation with phytoplankton carbon biomass on the northeastern Mediterranean Sea shore, with special emphasis on nanophytoplankton

**DOI:** 10.1101/745588

**Authors:** Merve Konucu, Elif Eker-Develi, Hasan Örek, Şehmuz Başduvar

**Affiliations:** Department of Biotechnology, Institute of Graduate Studies in Science, Mersin University, Ciftlikkoy, 33343 Mersin, Turkey; Faculty of Education, Mersin University, Çiftlikköy, 33343, Mersin, Turkey; Institute of Marine Sciences, Middle East Technical University, PO Box 28, Erdemli 33731, Mersin, Turkey

**Keywords:** Phytoplankton carbon biomass, marker pigments, CHEMTAX, nanoplankton, NE Mediterranean Sea

## Abstract

Summary Marker pigments are used as a proxy for biomass of distinct phytoplankton classes in different oceanic regions. However, sometimes disagreements are observed between microscopy and accessory-pigment based approaches in distinct regions mainly due to changing environmental factors governing diversity and structure of community composition. In this study, concordance between microscopy and HPLC-CHEMTAX methods were investigated first time in coastal waters of Erdemli, Turkey, in the Levantin Basin of the northeastern Mediterranean Sea by weekly intervals during 2015-2016. According to our results, marker pigment of diatoms, fucoxanthin, which was the most prominent pigment in the study area during most of the year, was a better indicator of diatom abundance than diatom carbon biomass. CHEMTAX derived values of diatom chlorophyll *a* (Chl *a*) were not in concert with either abundance or carbon biomass of this group. Contribution of dinoflagellates and cryptophytes to the phytoplankton community was underestimated with pigment based approach. Accessory pigment of cyanophytes, zeaxanthin, was also an important pigment in the samples. Biomass of haptophytes seemed to be overestimated by HPLC-CHEMTAX analysis. In contrast to diatoms, CHEMTAX derived chlorophyll *a* values of cryptophytes were correlated with abundance of this group but not with alloxanthin. Inclusion of live counts of nanoplanktic cryptophytes, haptophytes and prasinophytes provided a better correlation between microscopy and pigment based results. According to CHEMTAX analysis, nanoplankton and picoplankton constituted ∼55% of Chl *a* in the region.

## 1. Introduction

For quantification of standing stock of marine phytoplankton, cell counting, calculation of cell volume and carbon (C) or determination of chlorophyll *a* (Chl *a*) concentrations could be performed [1, 2, 3]. Among these parameters phytoplankton cell counting and calculation of cell volume is a process of extremely fatiguing and time consuming and not very suitable for analysis of high number of samples. For estimating standing stock from Chl *a* values obtained either through satellites or direct seawater measurements in a region, factors influencing C:Chl *a* ratios should be considered. This ratio varies between <10 and >200 among different phytoplankton classes as well as with change in nutrient concentrations, temperature, irradiance, growth phases and from species to species [3, 4, 5, 6, 7, 8]. Each of these parameters are variable in different seawater regions and should be validated with field data in less-studied locations. Chl *a* content of each phytoplankton group can be estimated based on their marker pigments determined by HPLC-CHEMTAX analysis and from these groups specific Chl *a* values, carbon biomass of phytoplankton groups could be assessed. HPLC analysis is faster than microscopy and provides one step analysis of all size groups through retention of pico-nano- and micro-phytoplankton on filters [9, 10]. However, verification of results obtained from pigment based approach by microscopy is a necessity at least for some of the samples due to sharing of some marker pigments by different phytoplankton classes, changing ratios of marker pigment:Chl *a* depending on species composition and growth phase, unusual pigment content of some dinoflagellates such as fucoxanthin (Fuco), alloxanthin (Allo), 19’-hexanoyloxyfucoxanthin (Hex), chlorphyll *b* (Chl *b*) [11, 12, 13, 14, 15].

In previous studies, HPLC-CHEMTAX method was generally found to be successful in estimating contribution of big-size diatoms and the main taxa to the total biomass but poor in estimation of small flagellates, haptophytes, prasinophytes and dinoflagellates [7, 8, 11, 16, 17, 18]. Small cells are especially important components of oligotrophic provinces in the oceans [19, 20, 21, 22, 23, 24]. Pico-cyanobacteria and nanoflagellates may contribute significantly to the total phytoplankton carbon biomass in coastal areas in different periods during seasonal cycle [23, 25, 26, 27]. These small sized organisms, especially the nanoplankton fraction, are generally disregarded with microscopy analyses due to difficulties in fixation, identification and counting processes of these motile and fragile groups [7, 23, 28]. There are previous studies in the region about picocyanobacteria abundance [29, 30] but species composition and abundance of nanophytoplankton is unknown in the study region especially for noncalcified haptophytes, prasinophytes and cryptophytes. There is only one previous investigation in a nearby region about marker pigments [24] but microscopy observation and CHEMTAX analysis are lacking in that investigation.

Our first goal was to determine first time the success of marker pigments to infer main phytoplankton taxa during a year by considering species diversity, abundance and carbon biomass in a coastal region having low productivity in the northeastern Levantine Basin of the Mediterranean Sea. The second aim was to observe the achievement of live counting process of nanoplanktic phytoplankton groups; haptophytes, prasinophytes and cryptophytes by comparing cell numbers with HPLC-CHEMTAX results.

## 1. Material and Methods

Samples were collected each week from the surface water of a pier on the Erdemli coast, Turkey (36°36’ N, 34°19’ E) in the north-eastern Mediterranean Sea during September 2015-September 2016 (Fig. 1).

**Figure 1.**
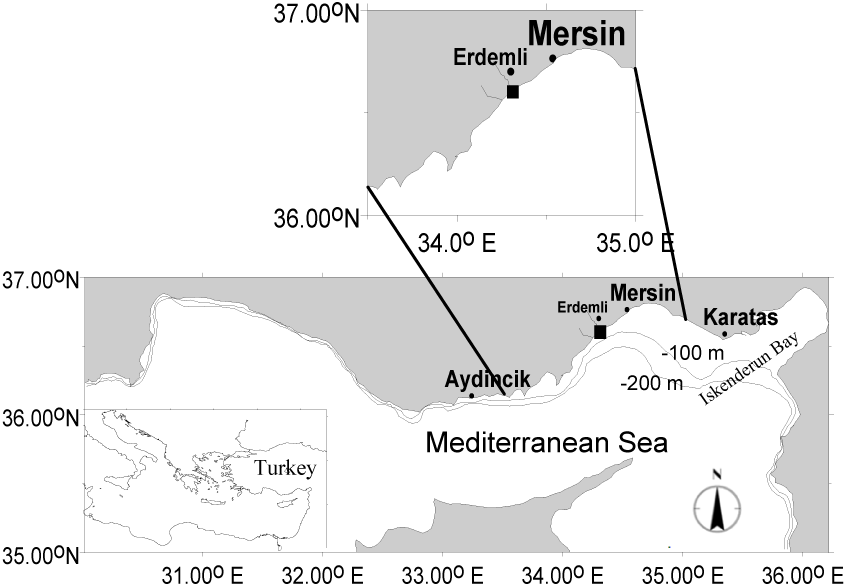
Sampling region.

### 2.1. General characteristics of the sampling region

Eastern Mediterranean is known as an ultra-oligotrophic sea mainly due to anti-estuarine circulation in the region [31, 32]. Average primary production of this region is 60-80 gCm^−2^y^−1^, which is about half of the production reported in other oligotrophic regions in the world’s oceans [32, 33, 34]. Average primary production value of a coastal station near to the sampling region was reported as ∼44 gCm^−2^y^−1^ [34]. Sampling region was reported to have good environmental conditions based on different Eutrophication Assessment Tools previously [35, 36]. External sources of nutrients in the sampling region are local rivers and the atmospheric dust coming from the Saharan and Arabian deserts (Appendix 1 in supplementary material, [3, 38, 39]. Eastern Mediterranean Sea has uniquely high N/P ratios (ranging from 25 to 28) compared to the west (22) and to the Redfield ratio (16) [32, 37, 40], which was attributed to high N/P ratios of all external inputs of nutrients to the eastern Mediterranean Sea, in addition to low denitrification rates [32]. The highest nitrate concentrations are recorded during winter-spring period at coastal waters [29, 39, 41]. Maximum flow rate of the Lamas River (∼7-12 m^3^/s) was reported between March-May [42]. Phytoplankton bloom period is recorded as February-early March in both coastal and open sea regions of the Mediterranean Sea related to winter mixing [38, 43, 44, 45]. Nutrients inflowing with this river sustain further phytoplankton growth during spring on coastal regions. Increases in nitrate concentrations are observed in summer months as well, but these concentrations do not reach to winter-spring levels near the sampling site [38, 39, 41]. The period between May and October is generally dry with very little precipitation in the sampling region [38, 46].

### 2.2. Phytoplankton sampling and analysis

Phytoplankton samples were collected from the surface into 1 L amber glass bottles and fixed with 31% formaldehyde buffered with borax to become 1.5% final concentration. Samples were settled 1-2 weeks and supernatant was siphoned by thin curved tubes. Phytoplankton cells (∼100-400 cells) were counted with a Sedgewick Rafter Cell under Nikon/eclipse TS100 inverted microscope. All area of the Sedgewick Rafter Cell was checked for rare species, a few columns were counted for abundant species. In addition to formaldehyde fixed samples (F), live cells (L) were also counted each week under microscope (Nikon/eclipse TS100) for small fragile cells of noncalcified haptophytes, cryptophytes and prasinophytes by putting direct seawater within a Petri dish within 1-3 hours following sampling. Certain amount of volume was checked each time by counting ∼50-200 cells depending on the density. Fine adjustment was carried out in each microscope field scanned for live cell counts. In order to understand possible mistakes originating from live counts, culture of a fragile, noncalcifying haptophyte species abundantly found in the region, *Chrysochromulina alifera* was counted both in live and fixed samples (0.2% final concentration of formaldehyde was used only for this sample and cells were counted immediately following fixation) [47, 48]. Abundance of *C. alifera* was 30% lower in live samples than in fixed ones. However, after a while fixed cells of *Chrysochromulina alifera* degraded in the samples. Cell counts of live and fixed prasinophytes, mainly *Pyramimonas* spp., were statistically correlated with each other (r^2^= 0.99, p<0.05), however, fixed cells were 5-6 times less in abundance or absent in fixed samples. Abundance of cryptophytes was also lower in formaldehyde fixed samples than in live samples and there was not a correlation between L-Crypto-C and F-Crypto-C (p>0.05). Thus, live counts of prasinophytes and cryptophytes were used for pigment and microscopy comparisons. For haptophytes, total carbon of live *Chrysochromulina* spp. and calcified haptophytes such as *Emiliania huxleyi* were used for comparison. Heterotrophic flagellates such as *Bodo* spp., *Metopian fluens, Ploeotia* spp., *Pteridomonas danica* were not included in live counts [49]. Photosynthetic cells of *Pyramimonas* spp., *Nephroselmis pyriformis, Pseudoscourfieldia* sp. and distinct cryptophyte species were counted as live. Some of the *Chrysochromulina* species in live counts could be mixotrophic.

Temperature and salinity were measured with a WTW LF330 model conductivity meter.

The volume (V) of each cell (100-1000 cells) was calculated by measuring its appropriate morphometric characteristics (i.e. diameter, length and width) [50, 51, 52]. One μm^3^ volume (V) was assumed equivalent to 1 pg wet weight [53, 54, 55]. Carbon biomasses were calculated from the volume of each cell throughout the text by the equations of Menden-Deuer and Lessard [56] as in [8].

### 2.3. HPLC pigment analysis

Known volumes (0.5 to 1 L) of seawater samples were filtered through Whatman 25 mm Ø GF/F filters under low vacuum and immediately stored at −20 °C deep freeze until the analysis within 3 months. Barlow et al. [57] method was adopted for the analysis. Pigment analysis was performed with an Agilent 1100 HPLC system. Samples were extracted with 90% HPLC grade acetone by using sonication. Extracted samples were kept overnight in a refrigerator and centrifuged before the measurements. Centrifuged samples were put into glass vials and placed inside of autosampler until injection. All injections were carried out by the autosampler. 200 μl of the extract was mixed with 200 μl 1 M ammonium acetate ion pairing solution by the autosampler [58]. Buffered extracts (100 μl) were injected through a 100 μl loop into a Thermo Hypersil MOS-2 C8 column (150×4.6mm, 3μm particle size, 120Å pore size and 6.5% carbon loading). Pigments were separated with linear gradient using a binary mobile phase system [24, 57]. Thirteen different phytoplankton pigments were detected by absorbance at 440 nm using an Agilent variable wavelength detector [58]. Pigment concentrations were calculated by ‘external standard’ equation [14]. The standards used were chlorophyll *a*, chlorophyll *b*, chlorophyll *c*2, peridinin, 19-butanoyloxyfucoxanthin, fucoxanthin, 19-hexanoyloxyfucoxanthin, diadinoxanthin, alloxanthin, lutein, zeaxanthin, divinyl chlorophyll-*a* and β-carotene (VKI, Denmark) (Table 2).

### 2.4. CHEMTAX processing

The CHEMTAX 1.95 program (excel version, [59, 60] was applied in order to obtain the taxonomic composition of phytoplankton from marker pigments. The 3 input matrices are: (1) the measured concentration of marker pigments and Chl *a* in the samples; (2) the theoretical input ratios of marker pigments to Chl *a* for each class of phytoplankton to be quantified; and (3) a ratio limit matrix restricting the iterative adjustments of these ratios operated by CHEMTAX. By using Matrix 1 and 2, CHEMTAX iteratively modify the difference between observed and calculated total pigment concentration utilizing changes in pigment ratios.

Eight phytoplankton groups; diatoms, dinoflagellates, haptophytes, cryptophytes, chlorophytes, prasinophytes, prochlorophytes and cyanophytes, were chosen for CHEMTAX analysis based on microscopy and pigment data. Pigments measured and their abbreviations are shown in Table 1.

**Table 1.** Abbreviations used in this article for phytoplankton groups, phytoplankton variables and photosynthetic pigments.

| 3 **Phytoplankton groups** | **Abbreviation** |
| --- | --- |
| 4 Diatoms | Diat |
| 5 Dinoflagellates | Dino |
| 6 Cyanophytes | Cyano |
| 7 *Prochlorococcus* sp. | Prochloro |
| 8 Chlorophytes | Chloro |
| 9 Prasinophytes in live samples | L-Prasino |
| 10 Prasinophytes in fixed samples | F-Prasino |
| 11 Calcified Haptophytes | Cal-Hapto |
| 12 Noncalcified Haptophytes in live samples | NonCal-Hapto |
| 13 Small flagellates | sFlag |
| 14 Cryptophytes in live samples | L-Crypto |
| 15 Cryptophytes in fixed samples | F-Crypto |
| 16 Raphidophytes, Silicoflagellates, Euglenophytes | Others |
| 17 |  |
| 18 **Carbon biomass** | -C |
| 19 **Abundance** | -A |
| 20 |  |
| 21 **Pigment** |  |
| 22 Chlorophyll *a* | Chl *a* |
| 23 Fucoxanthin | Fuco |
| 24 Chlorophyll *c2* | Chl *c2* |
| 25 Diadinoxanthin | Ddx |
| 26 Zeaxanthin | Zea |
| 27 19'-hexanoyloxyfucoxanthin | Hex |
| 28 Alloxanthin | Allo |
| 29 Chlorophyll *b* | Chl *b* |
| 30 Divinyl chlorophyll *a* | DVChl *a* |
| 31 Lutein | Lut |
| 32 Peridinin | Peri |
| 33 19'-butanoyloxyfucoxanthin | But |
| 34 β-carotene | Caro |

**Table 2.** Input [16, 59, 63] and output ratios of marker pigments to Chl *a* for the selected phytoplankton groups. Same input ratio matrix was used for high and low C:Chl *a* ratios. Abbreviations defined in Table 1.

|  | Fuco | Peri | Hex | Allo | Chl *b* | Zea | Chl *c2* | But | Lut | DVChl *a* | Chl *a* |
| --- | --- | --- | --- | --- | --- | --- | --- | --- | --- | --- | --- |
| **Input Ratios** |  |  |  |  |  |  |  |  |  |  |  |
| Diat | 0.81 | 0 | 0 | 0 | 0 | 0 | 0 | 0 | 0 | 0 | 1 |
| Dino | 0 | 0.547 | 0 | 0 | 0 | 0 | 0.28 | 0 | 0 | 0 | 1 |
| Hapto | 0.25 | 0 | 0.35 | 0 | 0 | 0 | 0 | 0.036 | 0 | 0 | 1 |
| Crypto | 0 | 0 | 0 | 0.35 | 0 | 0 | 0.13 | 0 | 0 | 0 | 1 |
| Chloro | 0 | 0 | 0 | 0 | 0.285 | 0.06 | 0 | 0 | 0.176 | 0 | 1 |
| Prasino | 0 | 0 | 0 | 0 | 0.623 | 0 | 0 | 0 | 0.035 | 0 | 1 |
| Cyano | 0 | 0 | 0 | 0 | 0 | 0.22 | 0 | 0 | 0 | 0 | 1 |
| Prochloro | 0 | 0 | 0 | 0 | 0 | 0 | 0 | 0 | 0 | 1 | 1 |
| **Output Ratios for high C:Chl *a* values ($\geq 8$)** |  |  |  |  |  |  |  |  |  |  |  |
| Diat | 0.81 | 0 | 0 | 0 | 0 | 0 | 0 | 0 | 0 | 0 | 1 |
| Dino | 0 | 0.547 | 0 | 0 | 0 | 0 | 0.28 | 0 | 0 | 0 | 1 |
| Cocco | 0.25 | 0 | 0.35 | 0 | 0 | 0 | 0 | 0.036 | 0 | 0 | 1 |
| Crypto | 0 | 0 | 0 | 0.35 | 0 | 0 | 0.13 | 0 | 0 | 0 | 1 |
| Chloro | 0 | 0 | 0 | 0 | 0.285 | 0.06 | 0 | 0 | 0.176 | 0 | 1 |
| Prasino | 0 | 0 | 0 | 0 | 0.623 | 0 | 0 | 0 | 0.035 | 0 | 1 |
| Cyano | 0 | 0 | 0 | 0 | 0 | 0.270 | 0 | 0 | 0 | 0 | 1 |
| Prochloro | 0 | 0 | 0 | 0 | 0 | 0 | 0 | 0 | 0 | 1 | 1 |
| **Output Ratios for low C:Chl *a* values ($< 8$)** |  |  |  |  |  |  |  |  |  |  |  |
| Diat | 0.81 | 0 | 0 | 0 | 0 | 0 | 0 | 0 | 0 | 0 | 1 |
| Dino | 0 | 0.547 | 0 | 0 | 0 | 0 | 0.28 | 0 | 0 | 0 | 1 |
| Cocco | 0.25 | 0 | 0.35 | 0 | 0 | 0 | 0 | 0.036 | 0 | 0 | 1 |
| Crypto | 0 | 0 | 0 | 0.35 | 0 | 0 | 0.13 | 0 | 0 | 0 | 1 |
| Chloro | 0 | 0 | 0 | 0 | 0.285 | 0.06 | 0 | 0 | 0.187 | 0 | 1 |
| Prasino | 0 | 0 | 0 | 0 | 0.623 | 0 | 0 | 0 | 0.035 | 0 | 1 |
| Cyano | 0 | 0 | 0 | 0 | 0 | 0.433 | 0 | 0 | 0 | 0 | 1 |
| Prochloro | 0 | 0 | 0 | 0 | 0 | 0 | 0 | 0 | 0 | 1 | 1 |

Initial accessory pigment:Chl *a* ratio matrices were shown in Table 2. Dividing diatoms to two subgroups in CHEMTAX input ratio did not help Diat-Chl *a* to better correlate with Diat-C or Diat-A (Appendices 1, 2 available as Supplementary Material).

Except pigments of cyanophytes and cryptophytes, zeaxanthin and alloxanthin, pigment:Chl *a* ratios were reported to remain almost stable under different light intensities [9, 16, 61, 62]. Thus, same pigment:Chl *a* ratios were used in CHEMTAX analysis for samples having both high and low C:Chl *a* ratios here.

The input pigment:Chl *a* ratio matrices used here included Chl *c2* and But but excluded Ddx and Caro, which are shared by many phytoplankton groups [14]. Chl c2 rather than Peri was significantly correlated with Dino-C (p<0.05, r^2^=0.86), thus this pigment was included in the input ratio matrix.

For adjusting potential changes in pigment:Chl *a* ratios, data set was separated to two parts based on high and low C:Chl *a* ratios as threshold being 8 (see also Appendix 4b, on which whole data set was run in CHEMTAX analysis).

The parameters set for the calculations were as follows: ratio limits were set to 500, weighting was ‘bounded relative error by pigment’, iteration limit = 5000, epsilon limit = 0.0001, initial step size = 25, step ratio = 2, cutoff step = 3000, elements varied = 5, subiterations = 1, weight bound = 30 [59, 64].

Linear regression analysis was performed in order to assess the relationship between random variables (pigment and/or carbon concentrations).

### 2.4. Grouping the dataset

For evaluation of relationship between microscopy and pigment based approaches, the dataset was subdivided to two groups as high and low C:Chl *a* samples. The threshold for high and low C:Chl *a* ratios was assumed as 8 by arbitrarily checking carbon and Chl *a* values. This number is close to the average C:Chl *a* ratio for the whole sampling period (9 ± 16). This threshold was defined as 25 in a timeseries study performed in the English Channel [7]. However, maximum carbon concentrations recorded in their study were several folds higher than values observed in the present study. Even though it was reported that low C:Chl *a* values corresponded to winter months in the mentioned study, there were many data points having low ratios in summer and autumn months as well in their study [7].

## 3. Results

### 3.1. Physicochemical parameters

Variations in temperature and salinity are shown in Fig. 2. The highest temperature values were recorded in July-August and the lowest values were recorded in January. Low salinity values were generally observed during winter-spring and in some autumn months.

**Figure 2.**
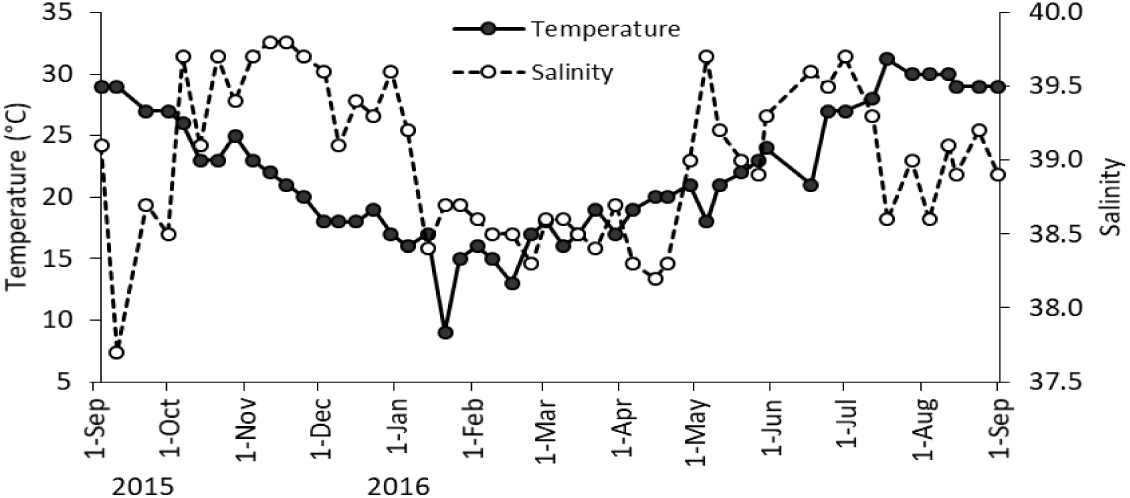
Variations in temperature and salinity during the sampling.

### 3.2. Annual averages of phytoplankton carbon biomass, abundance, marker pigments and CHEMTAX-derved Chl *a* values

According to annual average pigment concentrations, CHEMTAX assigned Chl *a* concentrations, phytoplankton carbon biomasses, diatoms were the most important group in the study area (Fig. 3). Although dinoflagellates was the second important group in terms of carbon biomass (Fig. 3d), unambiguous marker pigment of this group, Peri, was among the marker pigments having the minimum concentration. However, Chl *c*2 could be partially associated with dinoflagellates since there was a positive correlation between Dino-C and Chl *c*2 (r^2^ = 0.86, p<0.05). When CHEMTAX assigned Chl *a* values of phytoplankton groups and microscopy based carbon data were compared, contribution of haptophytes to the total Chl *a* seemed to be overestimated while the contribution of cryptophytes was underestimated (Fig. 3b and d). Carbon biomass of chlorophytes was higher than of prasinophytes due to a few big chlorophyte species such as *Halosphaera viridis* or some filamentous species (Fig. 3d). Based on HPLC results, Zea containing phytoplankton groups (i.e. cyanophytes, mainly picocyanobacteria including the genus *Prochlorococcus*) were also among the major contributors to the total pigment concentrations. According to CHEMTAX analysis cyanophytes were the second important phytoplankton group in terms of Chl *a* after diatoms among other groups (Fig. 3b). Unfortunately, biomass of this group (excluding filamentous ones) was not considered by light microscopy (Figs. 3d). Contribution of divinyl chlorophyll *a*, signature pigment of the cyanophyte *Prochlorococcus* sp., to the total marker pigment concentration was 3.2% only.

**Figure 3.**
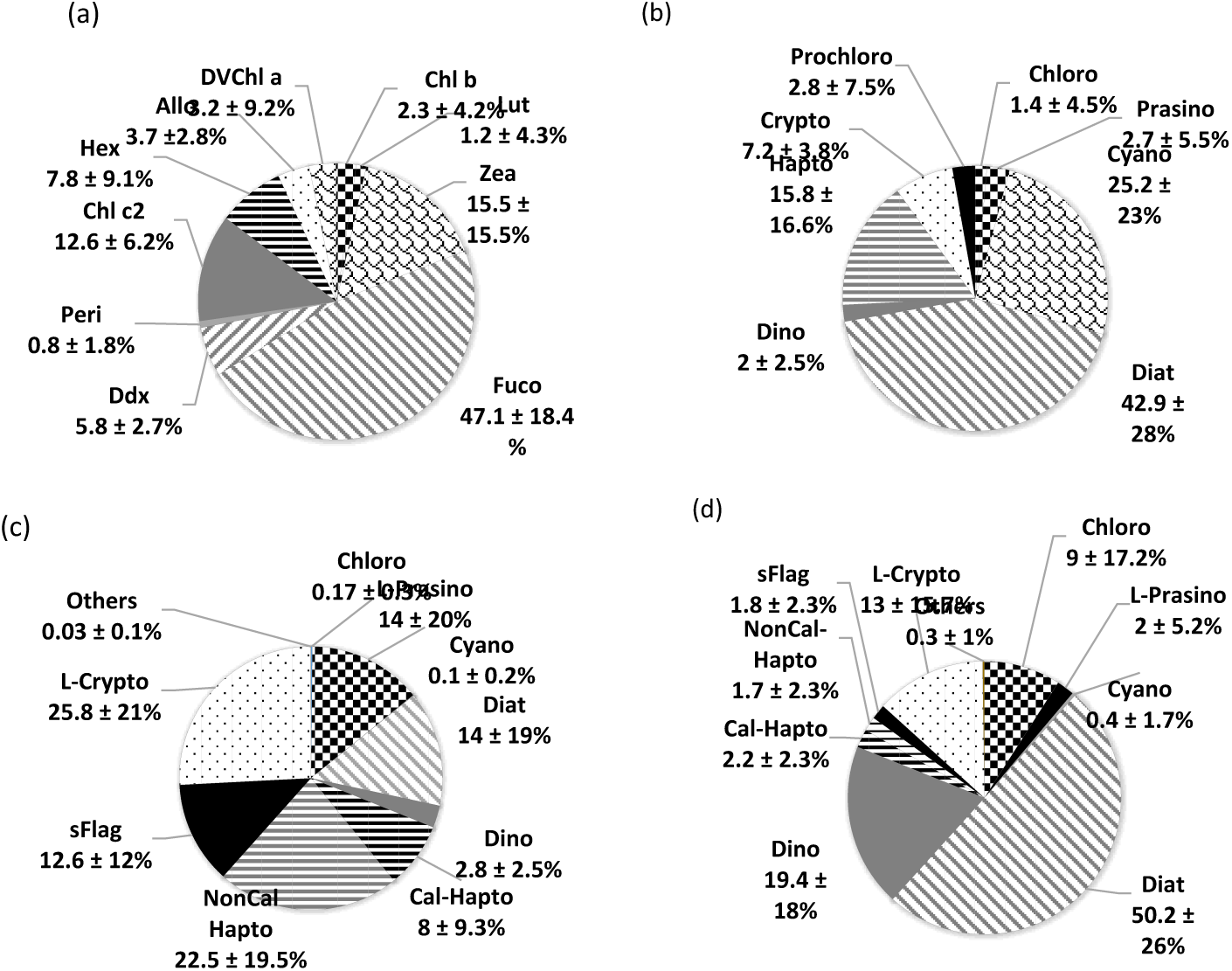
Annual average percentage contributions of (a) pigments to total accessory pigments (b) CHEMTAX derived phytoplankton groups to chlorophyll *a* (c) phytoplankton to total abundance and (d) phytoplankton to total carbon biomass during September 2015-September 2016 at the sampling location.

### 3.3. Weekly variations in phytoplankton carbon biomass, abundance, Chl *a* and C:Chl *a* ratios

Species number was quite high. A total of 222 species belonging to 15 different taxonomic classes were identified. 116 species out of 222 were diatoms. Main groups were shown in Table 1. Average Chl *a* concentration during the sampling was 1.19 ± 1.5 µg L^−1^ (n=50). Chl *a* values were <1 µg L^−1^ within half of the samples. When the whole dataset was taken into account, there was not any correlation between carbon and Chl *a* concentrations (p>0.05) during the sampling dates. However, when the samples were separated as high (≥8) and low (<8) C:Chl *a* ratios, a significant correlation in the high and low C:Chl *a* samples (r^2^=0.6, p<0.05, n=11 and r^2^=0.5, p<0.05, n=39, respectively) appeared (Fig. 4, pls see also Material and Methods, section 2.5). Since different factors such as light, temperature, species composition and nutrients control the variation of this ratio in different time periods, it is better to divide the dataset in timeseries studies. Average C:Chl *a* ratios were 32 ± 23 and 3 ± 1.6 in the high and low C:Chl *a* samples, respectively. The highest Chl *a* concentration observed on 3 February 2016, did not correspond to the highest carbon biomass during the sampling period (Fig. 4). Despite relatively low carbon biomass on 3 February 2016, Chl *a* reached to the highest level. High carbon values seen between January and May were mainly due to diatoms. The lowest phytoplankton abundance, carbon biomass and Chl *a* concentrations were observed in December-early January. Increase in Chl *a* concentration on 1 July 2016 was due to cyanophytes inferred from the rise in Zea concentration.

**Figure 4.**
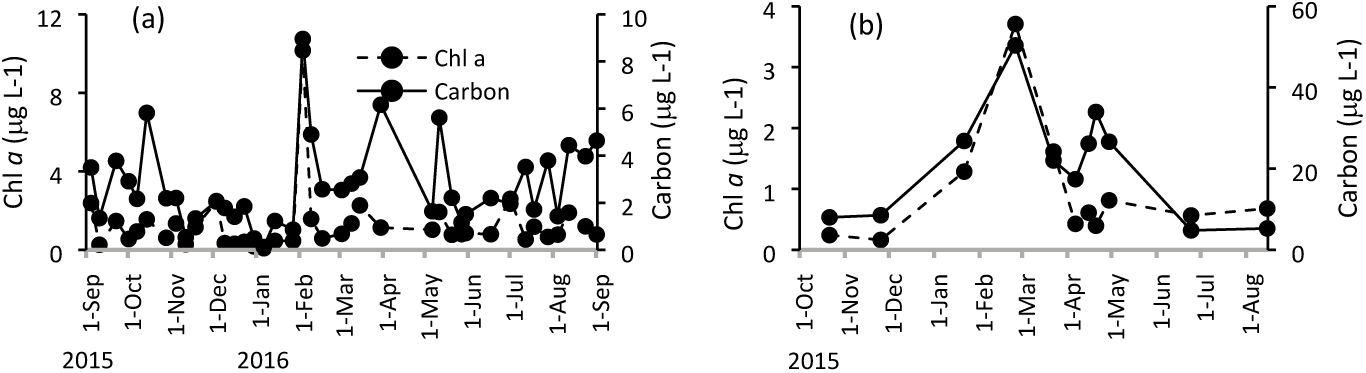
Variations in carbon and Chl *a* values in samples having (a) low C:Chl *a* ratios, <8 (b) high C:Chl *a* ratios, ≥8

### 3.4. Carbon biomass, abundance, marker pigments and Chl *a* of different phytoplankton groups

#### 3.4.1. Diatoms

Dominance of distinct diatom species in terms of carbon biomass on different dates has shown the interspecies competition (Fig. 5). Correlations between Diat-A and Fuco was much better than Diat-C and Diat-Chl *a* (Table 3, Fig. 6, Appendix 4a-d). Intraspecies variations in Fuco:Diat-C could be observed during 3 and 25 February 2016. *Asterionella glacialis* was dominant diatom species on both dates. This ratio was six times higher on the former date indicating exponential and stationary growth phases of this species on these dates, respectively. (Fig. 6).

**Table 3.** Relationship between Group-A-Group-C and marker pigments-Group-Chl *a* values in the samples having high (≥8) and low (<8) C:Chl *a* ratios. Cyanophytes were not correlated with Zea or DVChl *a* since picoplanktic cyanophytes were not counted.

|  | Low C:Chl *a* ratio |  | High C:Chl *a* ratio |  |
| --- | --- | --- | --- | --- |
| | $r^2$ | p | $r^2$ | p |
| Diat-C vs Diat-Chl *a* | 0.16 | <0.012 |  | >0.172 |
| Diat-C vs Fuco | 0.52 | <0.001 |  | >0.570 |
| Diat-A vs Diat-Chl *a* |  | >0.05 |  | >0.057 |
| Diat-A vs Fuco | 0.85 | <0.001 | 0.84 | <0.001 |
| L-Crypto-C vs Crypto-Chl *a* | 0.13 | <0.003 | 0.46 | <0.02 |
| L-Crypto-C vs Allo |  | >0.308 |  | >0.990 |
| L-Crypto-A vs Crypto-Chl *a* | 0.13 | <0.003 | 0.46 | <0.02 |
| L-Crypto-A vs Allo |  | >0.308 |  | >0.991 |
| T-Hapto-C vs Hapto-Chl *a* |  | >0.84 |  | >0.298 |
| T-Hapto-C vs Hex | 0.24 | <0.002 | 0.74 | <0.001 |
| Cal-Hapto-A vs Hapto-Chl *a* |  | >0.156 |  | >0.497 |
| Cal-Hapto-A vs Hex | 0.20 | <0.005 | 0.69 | <0.001 |
| L-Prasino-C vs Prasino-Chl *a* | 0.62 | <0.001 |  | >0.534 |
| L-Prasino-C vs Chl *b* | 0.39 | <0.001 |  | >0.542 |
| L-Prasino-A vs Prasino-Chl *a* |  | >0.762 |  | >0.534 |
| L-Prasino-A vs Chl *b* | 0.39 | <0.001 |  | >0.550 |
| Dino-C vs Dino-Chl *a* |  | >0.05 |  | >0.890 |
| Dino-C vs Peri |  | >0.942 |  | >0.893 |
| Dino-C vs Chl *c2* |  | >0.112 | 0.99 | <0.001 |
| Dino-A vs Dino-Chl *a* |  | >0.486 |  | >0.928 |
| Dino-A vs Peri |  | >0.508 |  | >0.758 |
| Dino-A vs Chl *c2* |  | >0.171 | 0.98 | <0.001 |

**Figure 5.**
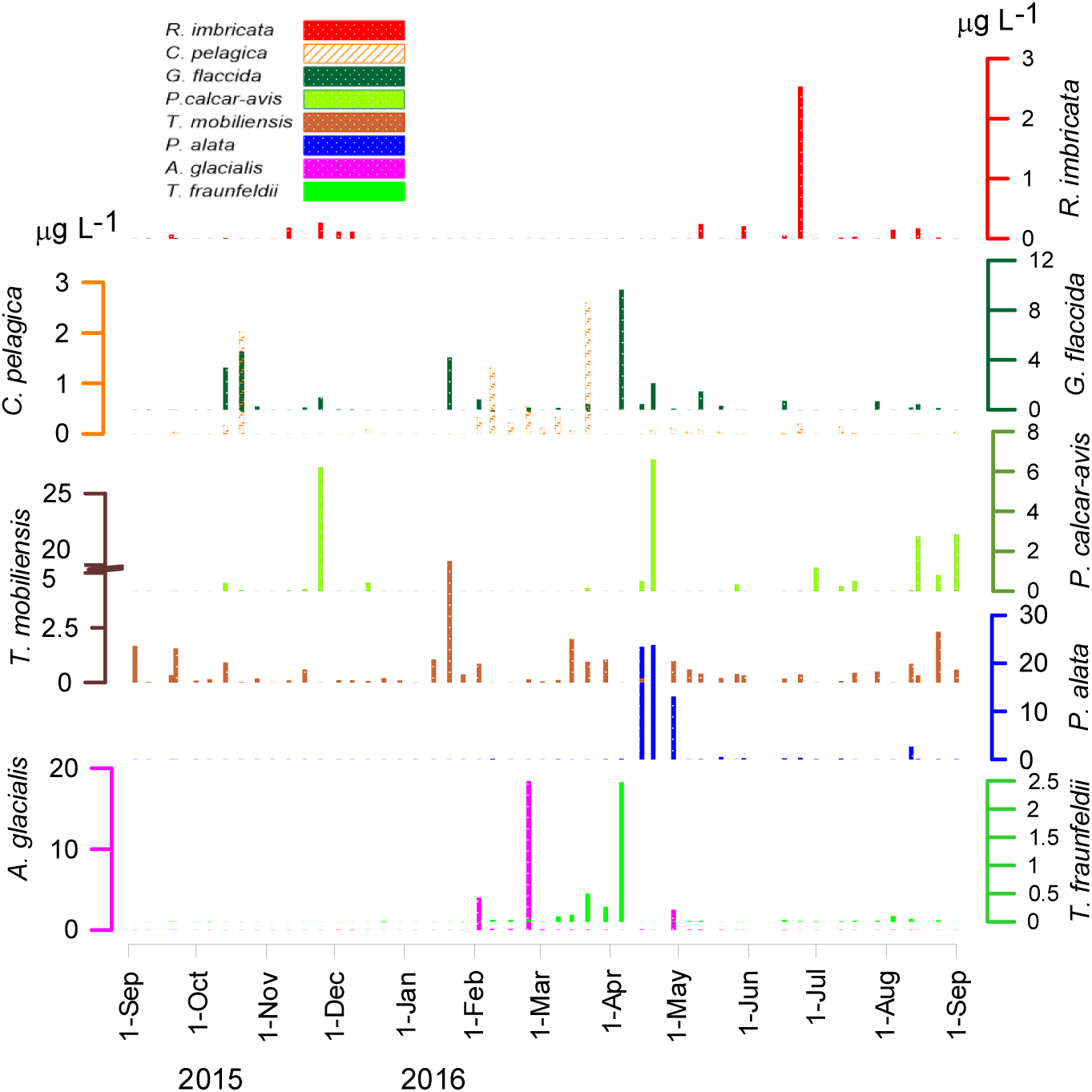
Diatom species having the highest carbon biomass values during 2015-2016.

**Figure 6.**
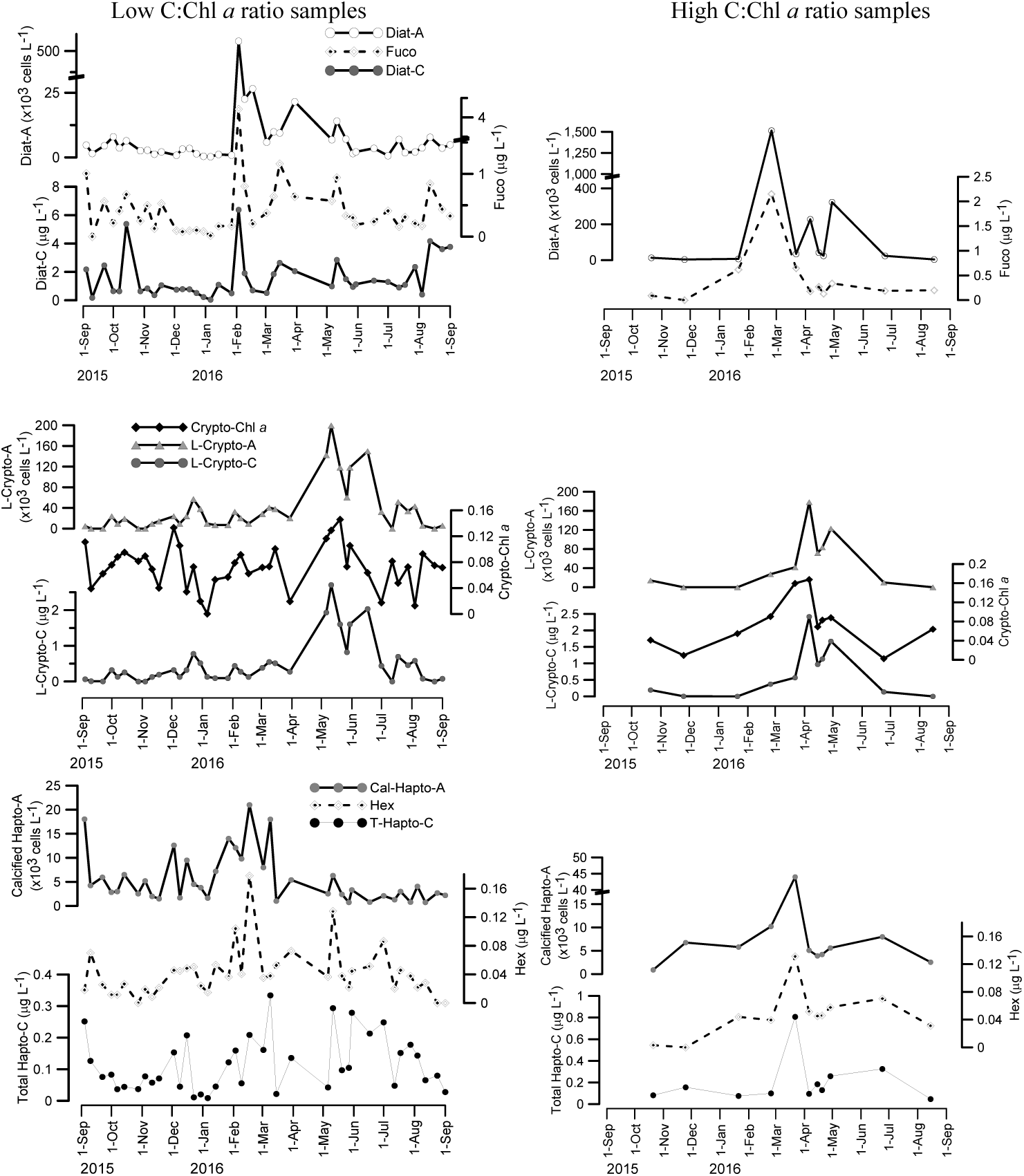
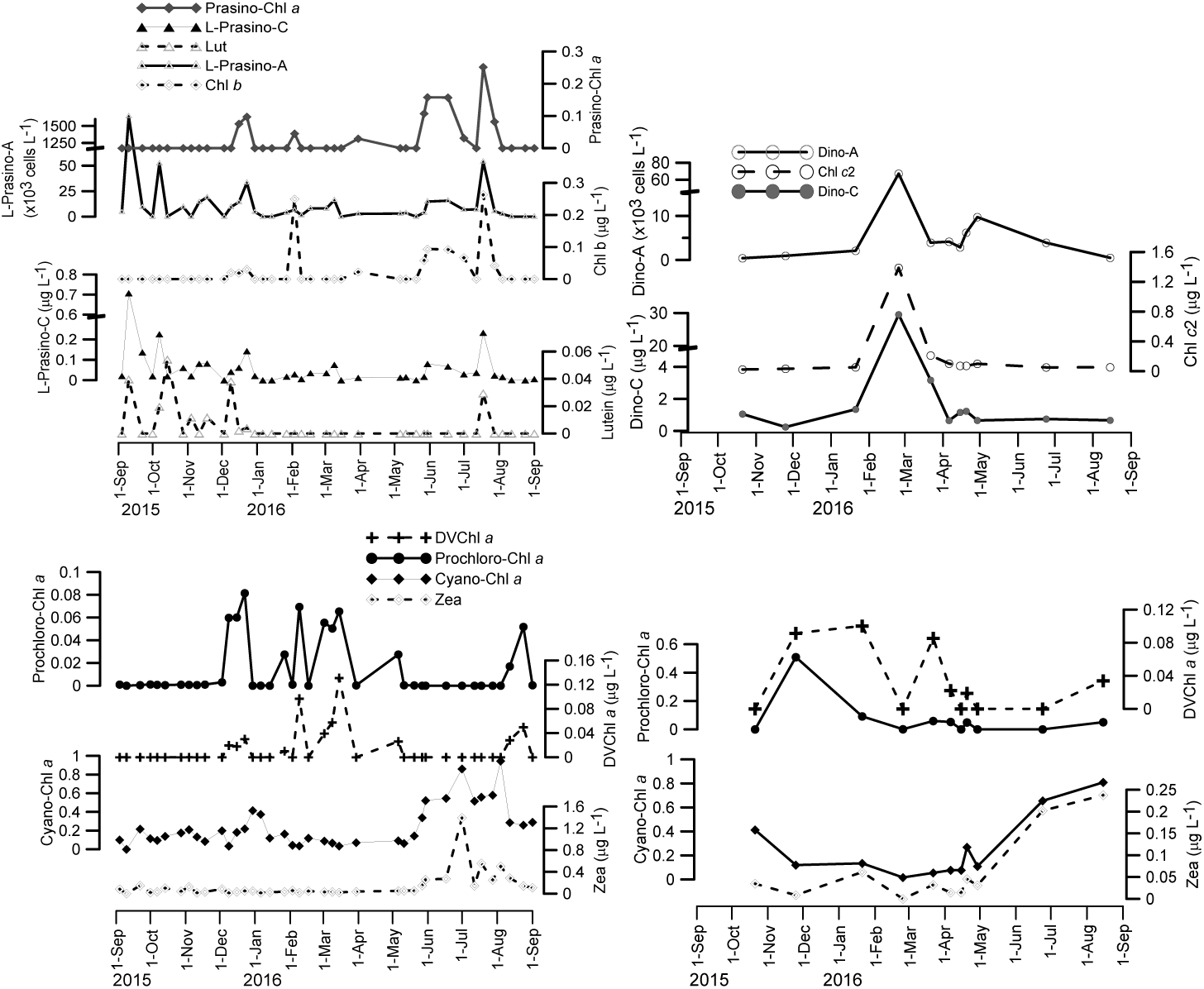
Variation in group carbon, cell abundance, marker pigment concentration and CHEMTAX assigned Chl *a* values of different phytoplankton taxa during September 2015-September 2016 in the sampling location when C:Chl *a* values were low (left side) and when C:Chl *a* values were high (right side). Pls see Table 2 for abbreviations. Only correlated parameters were shown.

Intraclass variations of Fuco:Diat-C were prominent during January, April and other months. Minimum values of Fuco:Diat-C were observed in January and April when big sized diatom species *Trieres mobiliensis* and *Proboscia alata* were dominant respectively (Fig. 5, 6). The highest Fuco:Diat-C ratio was observed on 3 February 2016 when relatively small-sized diatom *Asterionella glacialis* was dominant (Figs. 5, 6). There was not a correlation between CHEMTAX derived Diat-Chl *a* and Diat-C or Diat-A (Table 3). Dividing diatoms to two distinct groups in CHEMTAX input ratio did not provide a better correlation between Diat-Chl *a* and Diat-C or Diat-A (Appendix 3, supplemantary material).

#### 3.4.2. Cryptophytes

Cryptophytes were widespread during the sampling period in the study area. Abundance of this group was low in formaldehyde fixed samples while numerous individuals were counted in the live samples (Fig. 6). Annual average of Crypto-A in the live samples was ∼1000 times higher than the Crypto-A in the fixed samples. L-Crypto-A increased after March and remained high until July. Increase in Allo concentration during this period was not so conspicuous (pls see Appendix 4a). The highest Allo concentrations observed during February-March did not correspond to high Crypto-C values. Increases in Allo could be related to photoprotection, high pigment content of individuals during early growth phase or due to the ciliate *Myrionecta rubra*, which could be overlooked by microscopy. *Storeatula* cf. *major, Hemiselmis* sp., *Teleaulax* sp. and *Plagioselmis prolonga* were among the observed species. A few cells of *Myrionecta rubra* were recorded during May-August in live counts.

While there was a significant correlation between Crypto-C and CHEMTAX allocated Crypto-Chl *a*, Crypto-C and Allo were not correlated (p>0.05) (Table 3).

#### 3.4.3. Haptophytes

Although carbon biomass of haptophytes was less than chlorophytes and cryptophytes, marker pigment of this group, Hex had higher concentrations than Chl *b* and Allo values. There was a significant correlation between total Hapto-C and Hex values (Table 3). However, Hapto-Chl *a* values were not correlated with either total Hapto-A or total Hapto-C. While calcified haptophytes, mainly *E. huxleyi* was dominant during winter-spring period, noncalcified haptophytes, mainly *Chrysochromulina* spp., were abundant during spring-summer. As opposed to abundance values, carbon values of calcified haptophytes were higher than noncalcified haptophytes (Fig. 3c and d). Hex concentrations were higher when calcified haptophytes were dominant.

#### 3.4.4. Dinoflagellates

There was not any correlation between peridinin concentrations or Dino-Chl *a* and abundance or carbon biomass of dinoflagellates (Table 3). However, Dino-C was significantly correlated with Chl *c*2 in the high C:Chl *a* samples (r^2^ = 0.86, p<0.05, Fig. 6). Dino-C was generally lower than Diat-C but higher than carbon biomass of other nanoplanktic group, cryptophytes, haptophytes and prasinophytes (Fig. 3d). Major peak in dinoflagellate carbon biomass was on 25 February 2016. The dominant phytoplankton species on this date was *Alexandrium* sp.. Other dominant species during the year were *Gonyaulax* spp., *Protoperidinium* spp. and heterotrophic *Gyrodinium spirale*.

#### 3.4.5. Chlorophytes and Prasinophytes

Chlorophytes were generally represented by the subphylum Prasinophytina (referred as prasinophytes in the text) in the region rather than Chlorophytina. A few number of filamentous green algae and occasionally *Halosphaera* sp. were observed under the subphylum Chlorophytina. Majority of prasinophytes was composed of the genera *Pyramimonas, Nephroselmis pyriformis* and *Pseudoscourfieldia* sp. and flagellated stage of *Pterosperma* sp. was occasionally observed.

Prasinophytes reached to high abundances when diatoms and dinoflagellates were low in abundance. This period corresponded to low C:Chl *a* ratios and the correlation between Prasino-C and Prasino-Chl *a* was significant in this period, while no correlation was observed during high C:Chl *a* ratio period (Table 3, Fig. 6). During bloom concentration of the prasinophyte *Pyramimonas* spp. on 10 September 2015, Lut concentration was relatively high, while Chl *b* could not be detected (Fig. 6).

High values of Chloro-C in fixed samples were due to a few big chlorophyte species. Abundance of prasinophytes in live samples were 5-6 times higher than in fixed samples when present in both samples (e.g. 10 September 2015 and 22 March 2016).

#### 3.4.6. Cyanophytes

Picocyanobacteria was not included in the cell counts and there was not any correlation between abundance of fixed filamentous cyanophytes and Zeaxanthin concentrations (>0.05). Marker pigment Zeaxanthin has demonstrated that cyanophytes increased at the end of May and remained high until the beginning of August (Fig. 6). Zea was an important component of accessory pigments composing 14% of annual average accessory pigments. CHEMTAX allocated Cyano-Chl *a* constituted 25% of average Chl *a*.

DVChl *a* containing cyanophyte *Prochlorococcus* sp. was separately shown as Phrochloro-Chl *a* in CHEMTAX analysis. They reached peak values during winter-spring period (Fig. 6).

## 4. Discussion

### 4.1. Relationship between microscopy and HPLC-CHEMTAX approaches

According to our literature survey, HPLC-CHEMTAX results were not throughly compared with carbon biomass of phytoplankton groups in the Mediterranean Sea with a time series investigation. Previous studies focuses on impact of nutrients on phytoplankton composition and abundance mainly based on CHEMTAX results disregarding phytoplankton carbon biomass [24, 65, 66, 67]. Our results show that CHEMTAX assigned phytoplankton groups should not be assumed as a proxy to abundance or carbon biomass of diatoms, dinoflagellates and haptophytes in the study region located in the northeastern Mediterranean coast. Diatoms were the most important taxonomic class during majority of the sampling period which was shown by both microscopy and pigment data in the present study. However, CHEMTAX allocated Chl *a* values of diatoms were not correlated with carbon biomass of this group in general even if the data was separated into two parts as high and low C:Chl *a* ratio samples. Instead, direct concentrations of fucoxanthin were significantly correlated with abundances of diatoms during the sampling period (Table 3). High number of diatom species most probably having distinct marker pigment:carbon ratios (pls see section 3.5.1 in results) and variations in their abundance related to changing nutrient, light and temperature levels seem to complicate estimation of diatom carbon biomass via CHEMTAX analysis in the study region. Low fucoxanthin content of big-sized diatoms could easily be inferred from differences in peak levels of diatom carbon biomass and fucoxanthin maxima during the study especially in April (Figs. 6, Appendix 4a, c).

Second important group in terms of CHEMTAX analysis following diatoms was cyanophytes especially during warm period (Fig. 3b, Appendix 4b). Although carbon biomass of haptophytes was low, they were among major components of pigments and CHEMTAX derived Chl *a* following cyanophytes. Similar to diatoms, marker pigment of haptophytes, rather than group specific Chl *a*, were correlated with carbon biomass of this group. This could also be attributed to species diversity of haptophytes. The highest pigment diversity was found among haptophytes represented by five taxonomic groups in a study performed in the Nervión River estuary, Spain [69]. In contrast to diatoms and haptophytes, CHEMTAX derived Chl *a* values of cryptophytes and prasinophytes were correlated with carbon biomasses obtained from microscopy for these groups (Table 3). However, carbon biomass of cryptophytes appeared to be underestimated with HPLC-CHEMTAX based approach. Similar to the present study, CHEMTAX seemed to underestimate cryptophyte biomass but overestimate haptophyte contribution when compared to carbon values obtained from microscopy in a mesocosm study [17]. In a long term study performed in the western Mediterranean Sea during 2000-2014, major groups contributing to Chl *a* allocated by CHEMTAX were reported as haptophytes, diatoms prasinophytes and cryptophytes, respectively [67]. However, microscopy results were not presented in their study. Thus, how much haptophytes are important in terms of carbon biomass among other phytoplankton groups remains obscure. There was not any correlation between microscopy and HPLC-CHEMTAX results for dinoflagellates in the present investigation similar to previous studies [11, 69, 70].

It was striking that taking into account live counts of prasinophytes, cryptophytes and noncalcified haptophytes rather than fixed cells under light microscope provided a better correlation with pigment based approach in the present study. Abundance of these groups are either underestimated or cannot be differentiated taxonomically within fixed samples. 96.4% of phytoflagellates could not be attributed to any certain taxonomic group in fixed samples in a study performed in the southern Adriatic Sea [23]. Probably because of this, a weak correlation is found between pigment based and microscopy based results for some of these small-size phytoplankton groups reported in several distinct studies [7, 8, 16, 52, 71]. These small and mainly nanoplanktic groups dominated in the sampling region during warm period (April-August) similar to other investigations performed in the Mediterranean Sea [28, 68, 72,].

In this study, average contribution of Cyano-Chl *a* attributed by CHEMTAX to the total accessory pigments was estimated as 28% (Prochloro-Chl *a* + Cyano-Chl *a*, Fig. 3b). In the Adriatic Sea, picophytoplankton group was the most important fraction forming 49% of average biomass [23]. The highest Chl *a* values of cyanobacteria derived by CHEMTAX were observed during July and August in the present study which was consistent with flowcytometer results in a nearby location [29]. Average contribution of mainly nanoplanktic groups, cryptophytes, prasinophytes and haptophytes based on CHEMTAX was 25.7%. Based on microscopy, the share of nanoplankton fraction (haptophytes, prasinophytes, cryptophytes and small flagellates) within total carbon biomass and abundance were 21.4% and 83%, respectively, in the present investigation.

Total phytoplankton biomass and Chl *a* changed between <1 and 50 µg C L^−1^ and 0.1-10 µg L^−1^ in this study. Similar carbon values were observed during December 2000-February 2002 in a nearby sampling station [73]. Total phytoplankton biomass in the southern Adriatic Sea was between 8.5 and 80.7 µg C L^−1^ during 2006-2008 [23]. Carbon values found in the present study are lower than maximum values found in the Baltic Sea (965 µg C L^−1^, [52]) and in the Black Sea (∼800 µg C L^−1^, [8]).

### 4.2. C:Chl *a* ratio

In majority of the samples, C:Chl *a* ratio (n = 39) was low (average of 3 ± 1.6). When high ratios were grouped, the average became 32 ± 23 (n = 11). Our C:Chl *a* ratios were at the lower end of values previously reported in different geographic locations [8, 22, 52, 74, 75]. C:Chl *a* ratios based on fixed carbon values obtained from the results of primary production and total Chl *a* were also lower than 10 at a coastal station near to the present sampling station during monthly sampling performed in 2010-2011 [30]. It is known that Chl *a* content of phytoplankton increases under nutrient replete conditions compared to nutrient depleted ones. Thus, C:Chl *a* ratios increase under low nutrient conditions [4, 16, 38]. Average nutrient concentrations measured by the Middle East Technical University, Institute of Marine Sciences near to our sampling region (36°34’ N 34°16’ E) were relatively low during the same period with the present study in monthly samples; PO_4_-P: 0.05 µM (range 0.03 - 0.1 µM), NO_3_+NO_2_: 0.39 µM (range 0.13-0.84 µM), NO_2_–N: 0.06 µM (range 0.02-0.14 µM), NH_4_: 0.66 µM (range 0.21-1.26 µM) and Si 1.89 µM (1.11-3.18 µM). Consequently, low nutrient concentrations cannot explain generally low C:Chl *a* ratios in the region. The highest C:Chl a ratios were observed when big-sized diatoms were observed in April. Variations in C:Chl *a* ratios seemed to be related to prevalent species and their growth phases in distinct periods.

## 5. Conclusions

Fucoxanthin and 19’hexanoyloxyfucoxanthin were shown as better parameters for estimating abundances of diatoms and haptophytes than CHEMTAX allocated Chl *a* results in the study region. Carbon biomasses of cryptophytes and prasinophytes were correlated with CHEMTAX derived Chl *a* values for these groups. However, contributon of cryptophytes to total carbon biomass was much higher than its share in total Chl *a*. CHEMTAX results seem to overestimate contribution of haptophytes within total biomass based on our results similar to the investigation by Havskum et al. (2004).

Contribution of nano and picophytoplankton fraction to the Chl *a* was more than 54% in the present investigation. We conclude that CHEMTAX was unsuccessful in estimation of abundance or carbon biomass of the major phytoplankton group diatoms in the northeastern Mediterranean Sea. We have shown first time that live counts of nanophytoplankton provided a better correlation between microscopy and pigment based approaches for this size group in the study region.

## Acknowledgement

We acknowledge IMS-METU for providing us their nutrient data and for Turkish State Meteorological Service for supplying precipitation data. This study was supported by TÜBİTAK 115Y767 and Mersin University BAP 2016-2-TP-2-1912 projects.

